# A hierarchical Bayesian model to investigate trade-offs between growth and reproduction in a long-lived plant

**DOI:** 10.1101/2021.01.26.428205

**Authors:** Valentin Journé, Julien Papaïx, Emily Walker, François Courbet, François Lefèvre, Sylvie Oddou-Muratorio, Hendrik Davi

**Author notes:** **Corresponding author:** Valentin Journé. INRAE LESSEM, 2 Rue de la Papeterie, 38402 St-Martin-d’Hères, France.

## Abstract

A trade-off between growth and fecundity, reflecting the inability of simultaneously investing in both functions when resources are limited, is a fundamental feature of life history theory. This particular trade-off is the result of evolutionary and environmental constrains shaping reproductive and growth traits, but it remains difficult to pinpoint in natural populations of long-lived plants. We developed a hierarchical Bayesian model to estimate the inter-individual correlation among growth and reproduction, using observations at individual level over several years combined with resource simulations from an ecophysiological-based model (CASTANEA). In the Bayesian model, the resource, simulated by CASTANEA and incorporated as a latent variable, is allocated to tree growth, reproductive buds initiation and fruit maturation. Then, we used individual random effects correlated among energetic sinks to investigate potential trade-offs. We applied this original approach to a Mediterranean coniferous tree, Atlas Cedar (*Cedrus atlantica*), at two contrasted levels of competition, high versus low density population. We found that trees initializing many reproductive buds had a higher growth. Moreover, a negative correlation was detected between growth and fruit survival during maturation. Finally, trees investing more resource to maturate fruits initiated less reproductive buds. The level of competition did not impact the sign of these three correlations, but changed the level of resource allocation: low density population favored growth whereas high density favored reproduction. The level of resource have an impact on individual strategies. This new modeling framework allowed us to detect various individual strategies of resource allocation to growth versus late-stage reproduction on the one hand, and to early-versus late-stage reproduction on the other hand. Moreover, the sign of the correlation between growth and reproductive traits depends on the stage of reproduction considered. Hence, we suggest that the investigation of potential trade-offs between growth and reproduction requires to integrate the dynamics of resource and sink’s phenology, from initiation to maturation of reproductive organs.

## 2 Introduction

Trade-offs between life-history traits are expected to be ubiquitous throughout the living world. Although the concept of trade-off is used in many disciplines, we will follow here the definition in evolutionary ecology proposed by Stearns (1989): “trade-offs represent the costs paid in the currency of fitness when a beneficial change in one trait is linked to a detrimental change in another”. In this sense, a trade-off corresponds to a negative correlation between two traits related to fitness, which are observed on a set of individuals, or at different life-stages for a given individual. In particular, trade-offs between traits may occur when the resources, i.e. energy and nutrients, are limited. In this case, trade-offs are the results of resource allocation to different sinks sharing a common pool of resources. Thus, some of the trade-offs observed within a population may result from the variation in allocation strategies among individuals and/or among life-stages. The inter-individual variability of allocation strategies, as any other phenotypic trait, is expected to be driven by a combination of environmental and genetic factors (Garland and Carter, 1994). The theoretical model of van Noordwijk and de Jong (1986) demonstrated that the patterns of resource acquisition versus resource allocation can change the sign of the correlation between two life history traits: when resource acquisition is highly variable among individuals while the fraction allocated to each life history traits is similar, the between-trait correlation is positive. In the reverse case, when resource allocation is highly variable among individuals, but not resource acquisition, the correlation becomes negative. To better track trade-offs, one solution is to account for the variability of resources acquisition and allocation between individuals together with the measurement of life history traits: this approach requires combining ecophysiology with population ecology (Olijnyk and Nelson, 2013).

Trade-offs between reproduction and growth-related life-history traits have received much attention (Bell, 1980; Lovett Doust, 1989), in particular in annual and perennial plants (Obeso, 2002; Thomas, 2011; Lauder *et al*., 2019). Most studies investigated trade-offs through the phenotypic correlations between growth and reproduction at individual or population level, and reported either positive, negative or no correlations (Sánchez-Humanes *et al*., 2011; Thomas, 2011; Wu *et al*., 2020). Four main hypotheses related to resource allocation have been advanced to explain such idiosyncratic patterns (Pulido *et al*., 2014): resource allocation to growth and reproduction can be either (i) based on a hierarchy, where the resource is first allocated to reproduction and then to growth (Wardlaw, 1990; Suzuki, 2001) or (ii) linked to different resource pools, so that the resource allocated to growth is independent from that allocated to reproduction (Cremer, 1992; Yasumura *et al*., 2006; Knops *et al*., 2007; Żywiec and Zielonka, 2013) or (iii) linked to a single resource pool, with a constant fraction of resource allocated to each sink (Despland and Houle, 1997; Pérez-Ramos *et al*., 2010; Berdanier and Clark, 2016; Lebourgeois *et al*., 2018) or, finally, (iv) linked to a single resource pool with competition for resources allocated to different sinks (Koenig and Knops, 1998; Martín *et al*., 2015; Lebourgeois *et al*., 2018). Only in cases (i) and (iv) the expected trade-off can be observed, while cases (ii) and (iii) can lead to positive or non significant correlation. Besides resource allocation schemes, climatic conditions can have contrasted direct effects on reproduction and growth and generate negative environmental correlation without functional trade-off, for instance when favorable conditions for growth are unfavorable for reproduction (Knops *et al*., 2007; Mund *et al*., 2020).

Trade-off can also occur between early and late stages of reproduction. This is well illustrated by the *Quercus sp* study by Knops *et al.* (2007), where the meaningful trade-off occur between current and future reproduction. Indeed, due to the high reproductive costs in trees, an important seed production in a given year can decrease the level of resource to invest in future reproduction, explaining why less seeds are produced the year after (Sala *et al*., 2012). This trade-off between current and future reproduction was confirmed by removal experiments, where removing fruits increase the number reproductive buds and increase fruit production in the following year (Elmqvist *et al*., 1991; Fox and Stevens, 1991; Fox, 1995; Santos-del Blanco and Climent, 2014).

Another important trade-off in monoecious plants may occur between male and female reproduction. Studies investigating the cost of reproduction usually neglect the male function and in particular the abortion of reproductive flowers. The major assumption supporting this simplification is that male function needs fewer resources than female function (Elmqvist *et al*., 1991; Obeso, 2002) and is not limiting for reproduction. However, this hypothesis may no longer hold under changing climate (Schermer *et al*., 2019). In trees, studies of the trade-off between male and female reproduction are rare due to the difficulty to measure male reproductive biomass. But studying reproduction only from the female point of view may underestimate the initial reproductive effort and lead to a wrong estimate of correlation between growth and reproductive traits (Knops *et al*., 2007; Knops and Koenig, 2012). Hence, two hypotheses are commonly used: male biomass is relatively constant across years whereas reproductive female biomass follows available resource (Knops and Koenig, 2012), or both male and female biomass co-vary through time (Houle, 2001).

The aim of this study is to detect the trade-off between growth and reproduction at different stages of the reproduction cycle and considering different functions (male and female) within a tree population. To tackle this objective, we developed a new and original methodological approach by combining two different models for resource acquisition and allocation, respectively. On the one hand, we used an ecophysiological model to simulate the acquisition of resource at the individual tree level (Net Primary Production, NPP). This ecophysiological model simulates climate effects through several processes such as tree photosynthesis, respiration and soil evaporation. On the other hand, we developed a hierarchical Bayesian model based on resource allocation processes, which simultaneously accounts for (i) the timing of growth and reproductive processes, and (ii) the variation among individuals and years in resource allocation and in the phenotypic gender, both considered as latent variables. This approach allows to estimate the inter-individual correlation between growth and reproduction while accounting for resource heterogeneity between trees.

## 3 Material and Methods

### 3.1 Hierarchical Bayesian model

We developed a Bayesian hierarchical model linking resource to three energetic sinks: growth, reproductive buds initiation and female cones survival. Potential correlations between these sinks are accounted for by correlated random effects. The model is composed of two layers (see Fig. 1 for a general overview): the process model which describes how resource is allocated to each sink, and the data model which describes the links between the process model and the observations. The observations consist in repeated counts of male and female organs to characterize the reproductive sink and repeated measure of size increment to characterize the growth sink. Description of model implementation and procedure is present in Appendix S1: Section S2. We presented prior parameters in Table S1, accuracy of the model in Table S2 and posterior distribution in Table S3.

**Figure 1:**
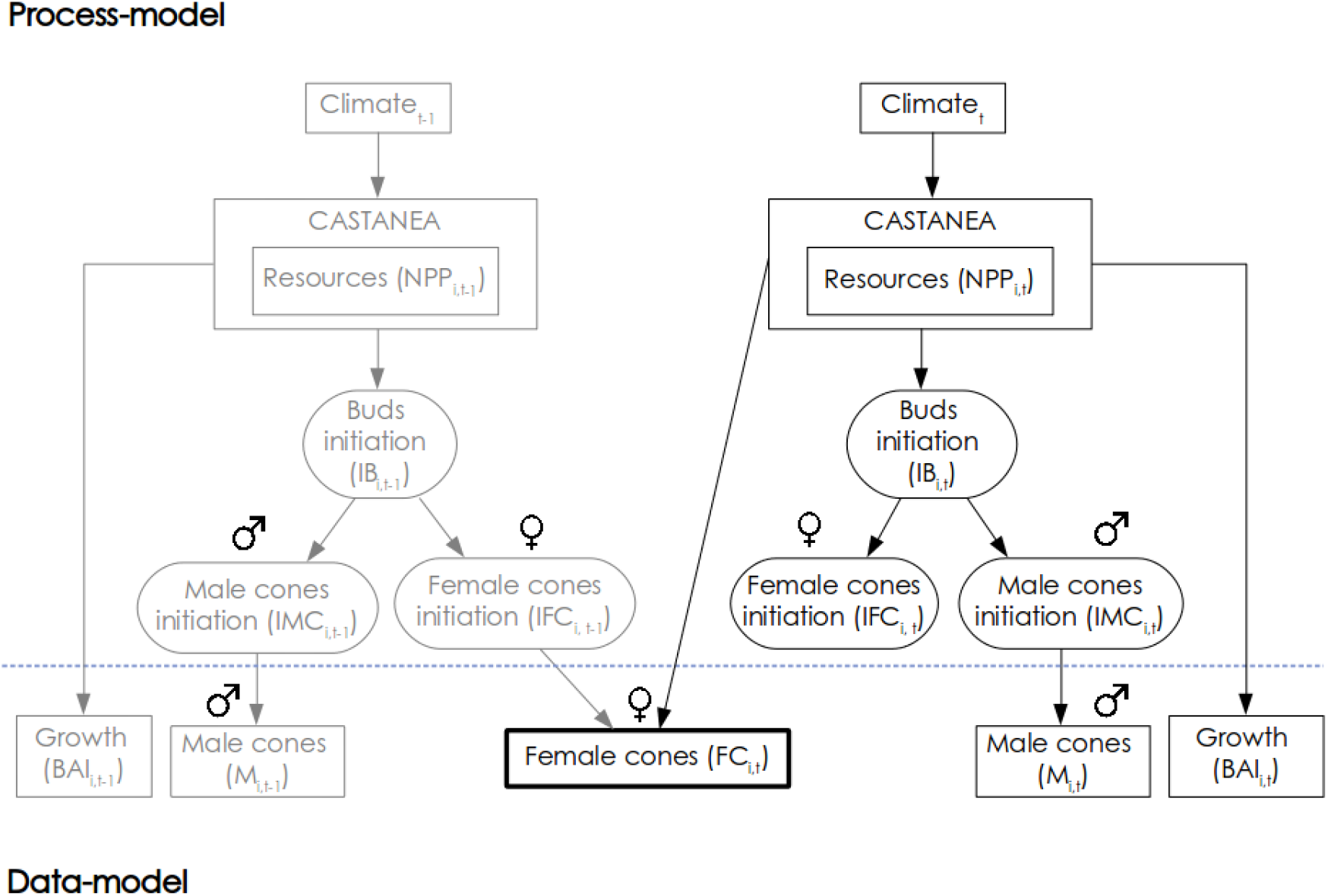
Graphical representation of the hierarchical Bayesian model used with the process and data models. Right angles boxes represent observed variables and elliptic boxes represent unobserved variable (i.e. latent variables). We represent the previous year (*t* −1) in gray color and current year (*t*) in black color. Resource (referred as *NPP*_*i,t*_) is simulated from the ecophysiological model (CASTANEA) with climate data. Resource is then allocated to radial growth (*BAI*_*i,t*_) and reproduction, which initiates buds (*IB*_*i,t*_). Buds differ then between male (*IMC*_*i,t*_) and females (*IFC*_*i,t*_) according to phenotypic gender (no presented in the figure, *PG*_*i,t*_). During the year *t* we have the maturation of males cones (*M*_*i,t*_) of the same year *t* and the the female cones survival based on female cones initiated the previous year *t* −1 (black bold box, *FC*_*i,t*_).

#### 3.1.1 Process model

##### Resource allocation to energetic sinks

The resource (*NPP*_*i,t*_) for individual *i* at year *t* determines its growth (*BAI*_*i,t*_), the number of initiated reproductive buds (*IB*_*i,t*_) and the probability of female cones survival 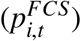. The use of correlated random effects (referred as *ϵ*_*x,i*_) allowed us to investigate potential trade-offs among these three energetic sinks. We tested two alternative models of inter-individual variation for each growth and reproduction trait to identify two potential types of phenotypic trade-off, either driven by NPP resource or not, respectively named “model 1” and “model 2”. In model 1, inter-individual variation acts on the capacity to valorize the amount of available resource (NPP), i.e. we introduced a random individual effect on the growth or reproduction trait response to NPP. In model 2, inter-individual variation is directly on the trait, i.e. we introduced a random effect on the intercept of the model. With this approach, there is no constraints on the triplet {*γ*; *β*_1_; *β*_2_}. Instead, the correlated individual random effect used here (*ϵ*_*x,i*_) allow more flexibility, and possible synergies or antagonisms between sinks to emerge from the model. To determine which one of the two model is the best, we included the variable *Y* ∼ Bernoulli (*p*_*Y*_) indicating whether NPP was the only resource responsible of inter-individual correlation (*Y* = 0) or not (*Y* = 1):

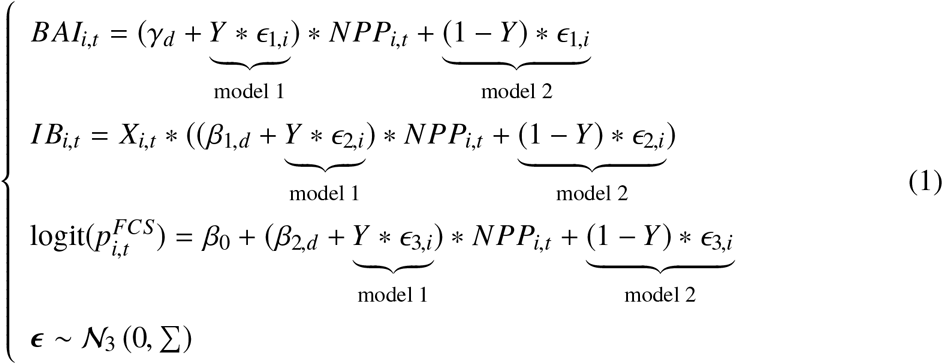

where *γ*_*d*_ is the slope parameter that depends on the density *d*. In the model 1, *ϵ*_1,*i*_ is an individual random effect associated to the slope parameter whereas in the model 2, the individual random effect does not constrains *NPP*_*i,t*_ on growth increment (*BAI*_*i,t*_). Then, *β*_1,*d*_ is the slope parameter than depends on density *d* and *ϵ*_2,*i*_ constrains the slope parameter (model 1) or is individual random effect for the effect of *NPP*_*i,t*_ on bud initiation (*IB*_*i,t*_) (model 2). Parameter *β*_2,*d*_ is the slope depending of density *d* and *ϵ*_3,*i*_ is associated to the slope parameter (model 1) or is the individual random effect for the effect of *NPP*_*i,t*_ on female cones survival 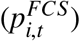 (model 2). *X*_*i,t*_is a Bernoulli variable (*X*_*i,t*_ ∼ ℬ (*p*_*X*_)), indicating if individual *i* produced (*X*_*i,t*_ = 1) or not (*X*_*i,t*_= 0) reproductive buds at year *t*, allowing to consider null values in the observations, because some trees never produced cones (see Fig 3). The intercept *β*_0_ is fixed to constrain 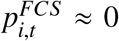 when *NPP*_*i,t*_ = 0. Note that the model does not account for other possible drivers of female cone survival such as pollen limitation.

**Figure 3:**
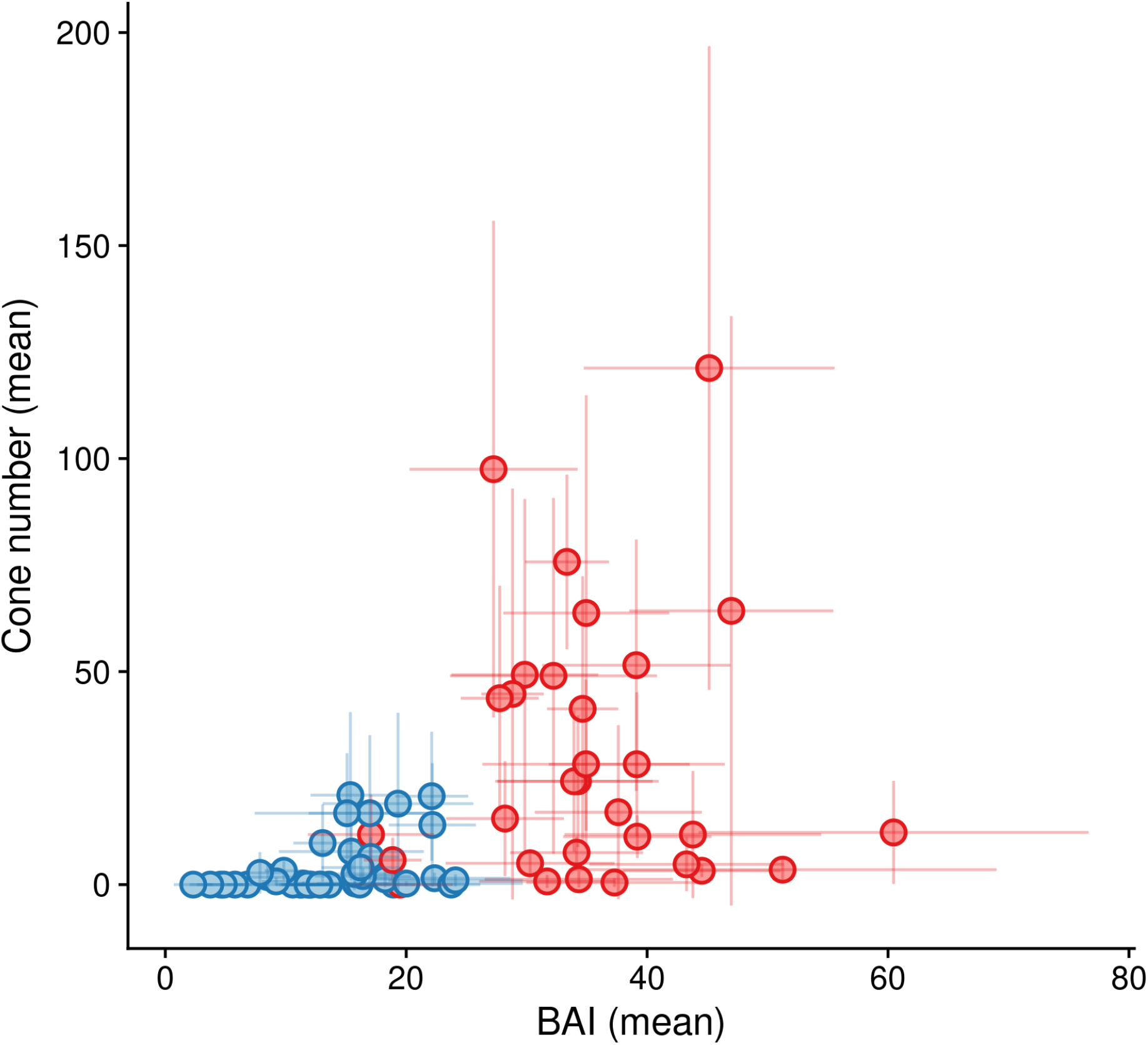
Raw data of female cone and growth for both density stand. Mean values with inter-annual standard deviations of female cone number 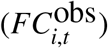 and mean growth increment 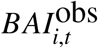, refereed here as BAI or Basal Area Increment in cm^2^ year^−1^) for each individuals. Blue is for high density (1200 stems ha^−1^) and red color for low density stand (250 stems ha^−1^).

This model explicitly considers that individual random effect on the three sinks, {*ϵ*_1,*i*_; *ϵ*_2,*i*_; *ϵ*_3,*i*_}, are related to each other through the variance-covariance matrix Σ. Their pairwise correlations can thus be used to investigate constrains in resource allocation to the three sinks. Indeed, for a given amount of resource available for an individual *i, NPP*_*i,t*_, the sign of the correlation between *ϵ*_*l,i*_, and *ϵ*_*k,i*_ indicates how much resource is respectively allocated to sinks *l* and *k*. Correlations were computed as *ρ*_*l,k*_ = Σ _*l,k*_/Σ _*l,l*_ Σ _*k,k*_).

##### Male and female reproduction

The model considers that initialized buds (*IB*_*i,t*_) develop into a number of initiated male cones (*IMC*_*i,t*_) and a number of initiated female cones (*IFC*_*i,t*_) according to the phenotypic gender (*PG*_*i,t*_) of individual *i* at year *t*, using the following model:

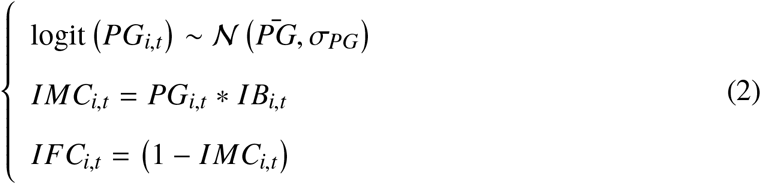

Here, phenotypic gender thus correspond to maleness, i.e. the ratio of male vs male and female initiated cones.

#### 3.1.2 Data model

The model uses repeated observations of growth, male reproduction and female reproduction, such as presented in our case in the section 3.3. We assume that observed growth 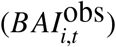 is related to the latent growth variable of the process model (*BAI*_*i,t*_) through:

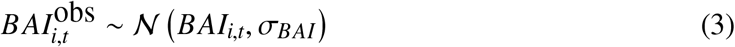

The number of initiated male cones (*IMC*_*i,t*_) is a continuous variable while the observed abundance of male cones 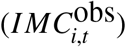 is a categorical ordered variable as described in section 3.3. To link *IMC*_*i,t*_ and 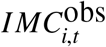 we used the following observational model:

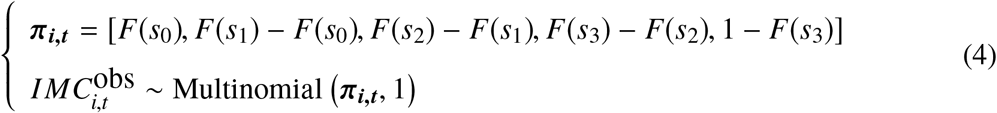

where *F*(.) denotes the cumulative distribution function of a normal distribution with mean *IMC*_*i,t*_ and variance σ_*IMC*_. {*s*_0_, *s*_1_, *s*_2_, *s*_3_} is a set of fixed thresholds determining the boundaries between each value of the notation, and are derived from *F*(.). Note that this approach is equivalent to consider a probit link in the case of binary data.

Finally, the observed count of female cones 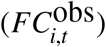 is linked to the latent variable describing number of current bud initiated 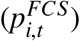 and previous year reproduction (bud initiation, *IB*_*i,t*−1_, female cones initiation, *IFC*_*i,t*−1_) through the following Poisson observational model:

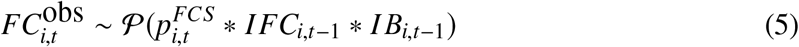

### 3.2 Study species and site

We applied this model to Atlas cedar, *Cedrus atlantica* (Manetti ex Endl.) Carrière, a Mediterranean coniferous tree species. This monoecious species reaches it sexual maturity between 15 and 30 years old (Toth, 1978), and carries male and female organs irregularly dispersed on the crown of the tree. Male reproduction, from reproductive bud initiation to pollen maturation, is achieved within one year, whereas female reproduction, from initiation to cone maturation, spreads over two years (Fig. 2). Reproductive buds are initiated during summer (June-July for male buds and late August for female buds) at year *t-1*, followed by pollination in September, when female cones open and receive pollen. Female cones close their scales in October-November, when male cones fall down. A time lag between pollination and fecundation characterizes coniferous species (Williams, 2009), with a duration of nine month for *C. atlantica*. Pollen germinates in spring at year *t*, then ovules are fertilized and seeds begin to maturate until autumn of the same year, and mature seeds are dispersed in year *t+1* (Toth, 1978). Hence, two generations of female cones may co-exist on a tree and it is possible to distinguish green one-year cones from brownish two-years cones. *Cedrus atlantica* can be referred as a masting species, characterized by a seed production highly variable among years and synchronized among individuals (Kelly and Sork, 2002; Krouchi *et al*., 2004).

**Figure 2:**
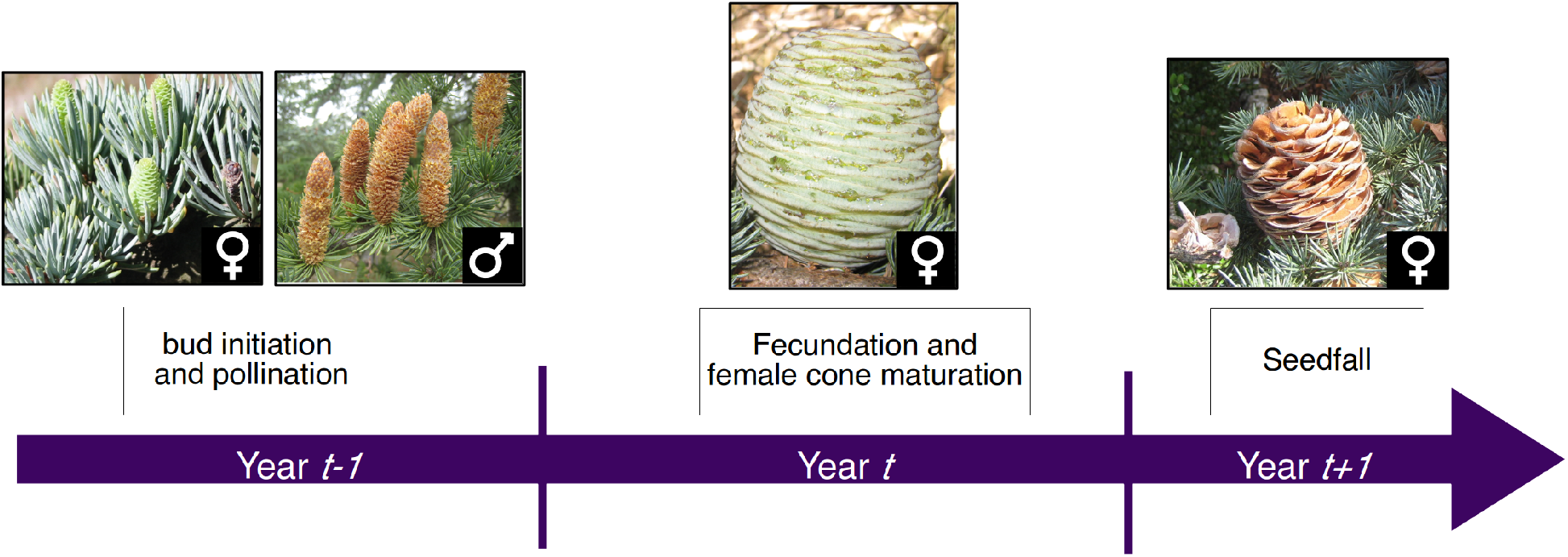
Overview of the general reproductive and growth cycle of *Cedrus atlantica*. Male reproduction and growth is carry on in one year. The duration of female reproduction takes two years from bud initiation during *year*_*t*−1_ followed by the maturation of female cone during *year*_*t*_. Seedfall occurs during *year*_*t*+1_.

The study site is a 35 years-old experimental plantation located in Mont-Ventoux in France, a Mediterranean mountain, at 1170 meters of elevation (44°07’ 05” N, 5°20’ 38” E). All the trees were planted in similar pedo-climatic conditions. The initial tree density was 2700 stems ha^−1^. In this experiment, two thinning strategies had been applied leading to contrasted competitor densities at the time of observations: high (1200 stems ha^−1^) versus low density (250 stems ha^−1^). More details on the silvicuture experiment and tree growth response are available in Guillemot *et al.* (2015).

### 3.3 Observations of growth and reproduction

We monitored 40 individual trees in the high-density stand and 31 individual trees in the low-density stand. These individuals were randomly sampled within each stand and measured each year from 2002 to 2005 (except for growth, with a longer dataset from 1989 to 2015). We first measured the diameter at 1.3 meters (*DBH*_*i,t*_) for each individual tree *i* and each year *t*. Annual basal area increment 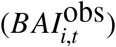 was computed as: 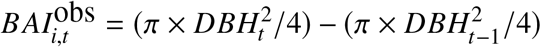 Male cones abundance (*M*_*i,t*_) was recorded as a qualitative ordered variable, consisting in a score ranging from 0 to 4: “0” means no male cone is observed; “1” means few male cones are dispersed in the canopy; “2” means male cones are abundant in one branch; “3” means male cones are abundant on two branches and “4” means male cones are abundant over the whole tree canopy. These scores were converted into multinomial observations 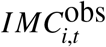 as follows:

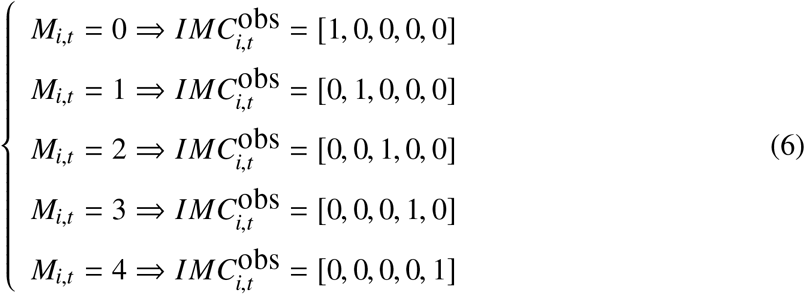

Female cones 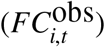 were individually counted over the whole canopy from the ground with binoculars and at two seasons each year (in spring then in late summer) to discriminate one-year versus two-year female cones based on their color. The identification and distinct count of one-year versus two-years cones allowed us to determine the number of cones produced each year. The raw relationship between 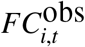 and 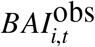 in Fig. 3 shows a positive correlation between growth and female reproduction and individual variation.

### 3.4 Resource simulation using CASTANEA

We used the ecophysiological model CASTANEA to estimate the carbon resource available each year for each tree. CASTANEA aims to simulate carbon and water fluxes of monospecific forest ecosystems (Dufrêne *et al*., 2005). The model simulates radiation transfer, photosynthesis, autotrophic respiration, carbon allocation to different tree compartments, evapotranspiration and water balance. A complete description of the model is presented in Dufrêne *et al.* (2005) with subsequent modifications described in Davi *et al.* (2009) and in Davi and Cailleret (2017). The model input data are daily climate data, soil characteristics (texture, depth and stone content), and initial tree characteristics (height, diameter and age). We calibrated and validated the model for *C. atlantica* on our study site for both stand densities. The values of CASTANEA parameters, simulations design, and validation procedure are described in Appendix S1: Section S1. In the following hierarchical Bayesian model, resource was modeled for each individual through the Net Primary Productivity (*NPP*_*i,t*_) simulated from 1999 to 2006, and corresponding to the difference between gross primary productivity and autotrophic respiration.

## 4 Results

### 4.1 Resource and phenotypic gender differences between densities

The model fitted accurately male fecundity (Brier score closed to 0). We obtained a lower accuracy for both female cones production (Bayesian p-value = 0.84) and tree growth (Bayesian p-value = 0.18). The posterior predictive checks of the hierarchical Bayesian model are presented in Appendix S1: Table S2. All trees had a high probability to be reproductive in both plots (*p*_*X*_, posterior median [CI95%] = 0.84 [0.79; 0.88]). Remaining posterior parameters are described in the following part and in Appendix S1: Table S3. The resource *NPP*_*i,t*_ simulated with CASTANEA model varied between years (due to climate) and between individuals (due to variations in tree size and stand density; Appendix S1: section S1). *NPP*_*i,t*_ had a mean value of 11990 gC tree^−1^year^−1^(sd = 3030 gC tree^−1^year^−1^) and 8751 gC tree^−1^ year^−1^ (sd = 2335 gC tree^−1^ year^−1^), respectively for the low-density and high-density plot (Fig. 4 a).

**Figure 4:**
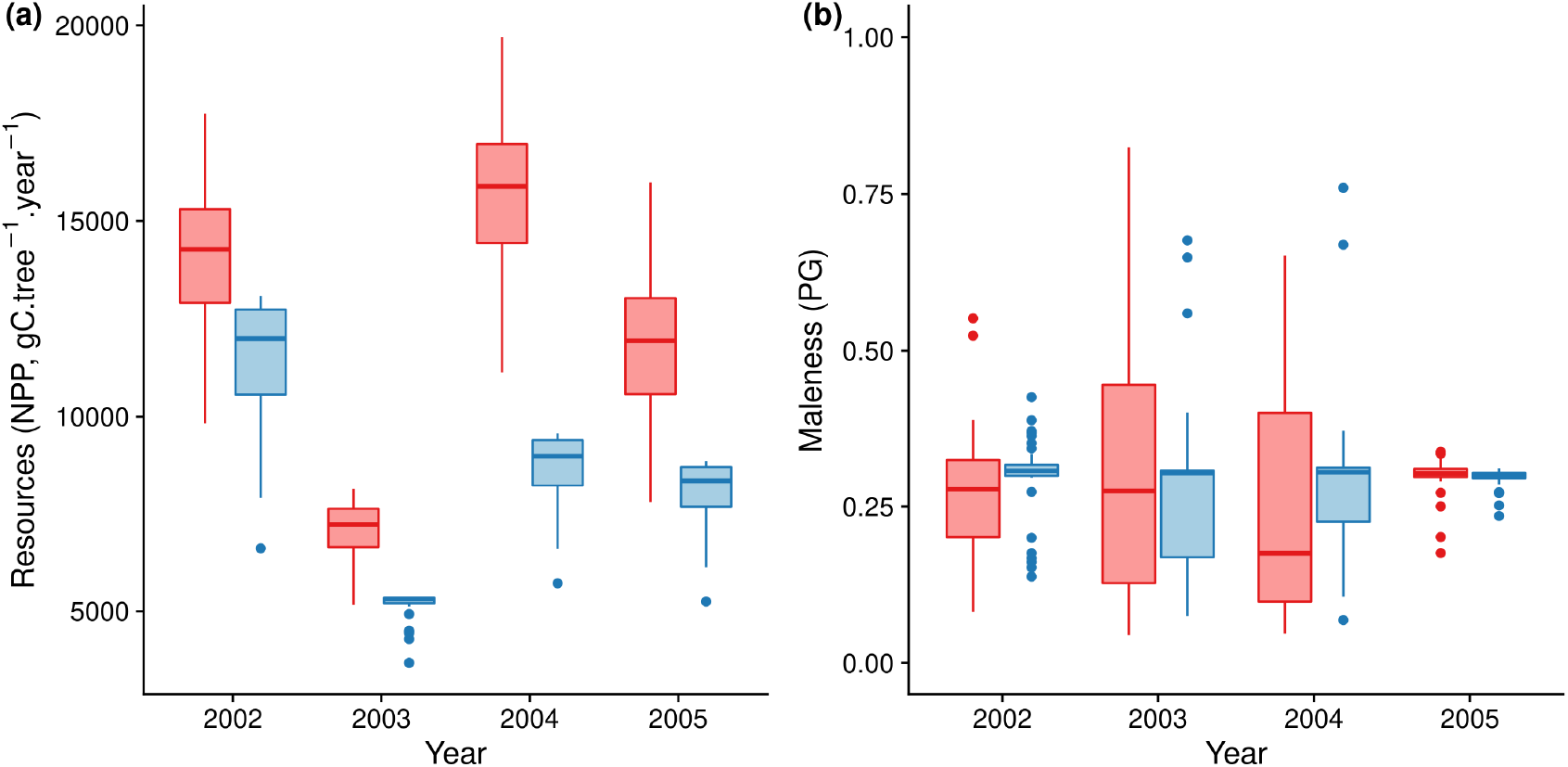
Output simulation of resource obtained from the ecophysiological model CASTANEA and estimation of the phenotypic gender from the hierarchical Bayesian model. (a) Boxplot of resource (Net Primary Productivity, in gC m^−2^ year^−1^) simulated with the CASTANEA model for both stand density and years (b) Boxplot of phenotypic gender estimated for both stand density and years, ranging in y-axis from femaleness (0) to maleness (1). For both graphics, blue color is for the high density (1200 stems ha^−1^) and red for the low density stand (250 stems ha^−1^).

The phenotypic gender did not differ between the two density stands, with a mean maleness equal to 0.29 (sd = 0.04) for the high-density stand and to 0.28 (sd = 0.06) for the low-density stand (Fig. 4 b).

### 4.2 Correlation between individual random effects

We found that Y was equal to 0 for all iterations, meaning that the best model to infer trade-offs in resource allocation is the model 2, where the *ϵ*_*x,i*_ are not linked to NPP. We reported the posterior distribution of correlations in Fig. 5 (a), (b) and (c) and pairwise covariations between individual random effects *ϵ*_1,*i*_; *ϵ*_2,*i*_; *ϵ*_3,*i*_ in Fig. 5 (d), (e) and (f).

**Figure 5:**
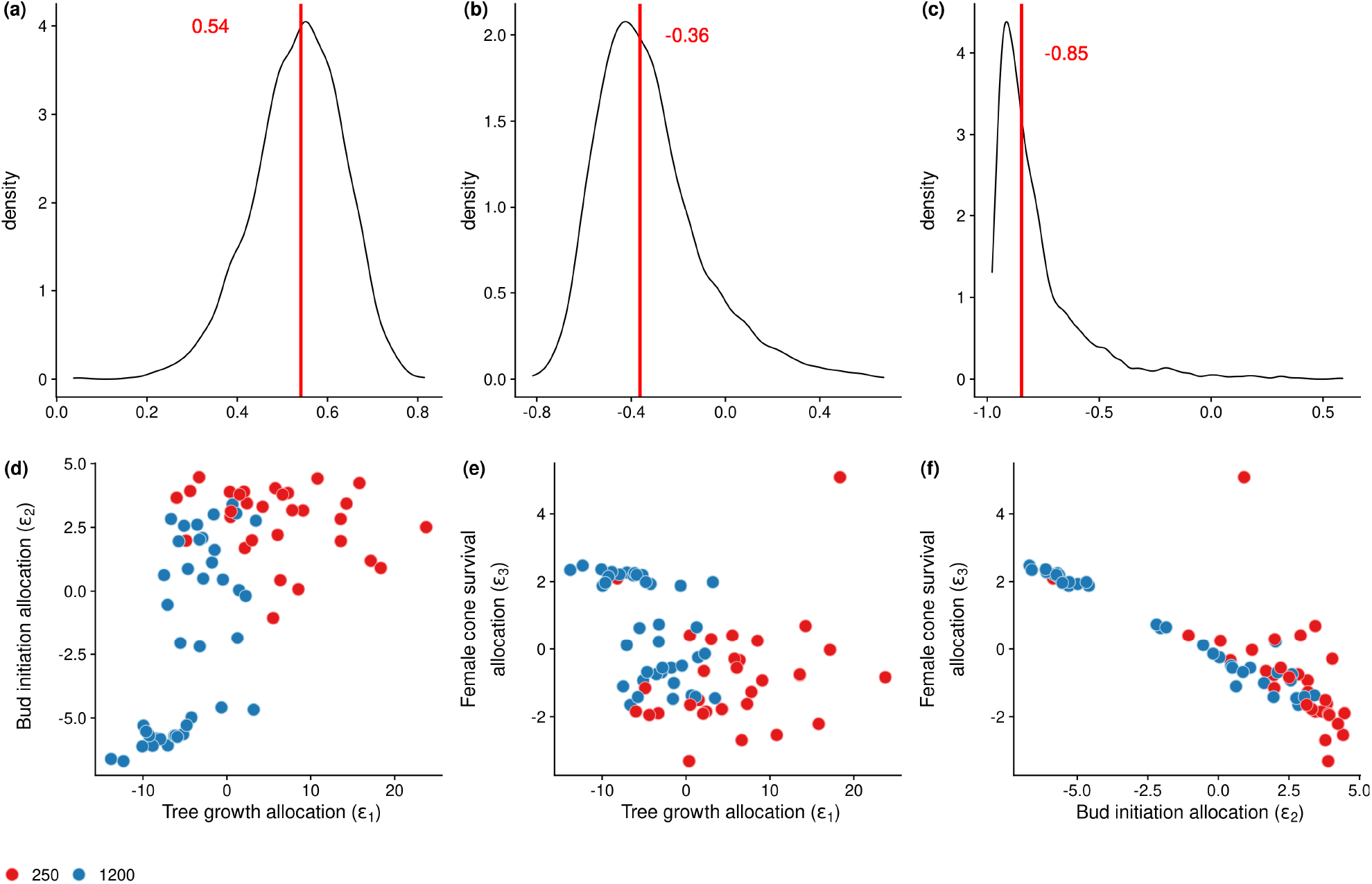
Correlation among sinks and their probabilities. (a) Density of the correlation *ρ*_1,2_ is positive (b) Density of the correlation *ρ*_1,3_ is negative (c) Density of the correlation *ρ*_1,2_ is negative. The red line represent the median posterior value for these three graphics. (d) Positive correlation between growth allocation (*ϵ*_1_) and allocation to buds initiation (*ϵ*_2_) (e) Negative correlation between growth allocation (*ϵ*_1_) and female cone survival (*ϵ*_3_) (f) Negative correlation between allocation to buds initiation (*ϵ*_2_) and female cone survival (*ϵ*_3_). Dots represent mean individual values, with blue and red color, respectively for high and low density of the stand.

The estimated correlation between inter-individual variations in growth (*ϵ*_1,*i*_) and reproductive buds initiation (*ϵ*_2,*i*_) was positive (*ρ*_1,2_ = 0.54[0.32, 0.70] with *pr*[*ρ*_1,2_ > 0] = 1, Fig. 5 a,d). Secondly, the estimated correlation between inter-individual variations in growth (*ϵ*_1,*i*_) and female cones survival (*ϵ*_3,*i*_) was significantly negative (*ρ*_1,3_ = −0.36[−0.65, 0.24] with *pr*[*ρ*_1,3_ < 0] = 0.91, Fig. 5 b,e) indicating a trade-off. Lastly, the estimated correlation between inter-individual variations in reproductive buds initiation (*ϵ*_2,*i*_) and female cones survival (*ϵ*_3,*i*_) was negative (*ρ*_2,3_ = −0.85[−0.96, −0.19], with *pr*[*ρ*_2,3_ < 0] = 0.98), indicating a clear phenotypic trade-off between bud initiation and female cone survival (Fig. 5 c,f). These three correlations mainly consist in between-density correlations and within-density correlation in the high density stand (low resources), but only one within-density correlation is found in the low density stand (high level of resources) between female cone survival and bud initiation.

### 4.3 Effects of stand density and phenotypic gender on individual allocation strategies

We found a higher growth response to available resource in the low density stand than in the high density stand (*pr*[*γ*_1200_ > *γ*_250_] = 0.25). However, we found a higher reproductive response, for initiation and maturation, to available resource in the high density than in the low density (*pr*[*β*_1,1200_ > *β*_1,250_] = 0.86 and *pr*[*β*_2,1200_ > *β*_2,250_] = 0.94). The ratio of resources allocated to growth was higher in the low density stand, whereas the ratio of resources allocated to reproduction was higher in the high density stand. Lastly, individual allocation strategies (*ϵ*_*x,i*_) were not affected by the phenotypic gender. Probabilities to obtain a significant correlation *P* < 0.05 between *ϵ*_1,*i*_ and phenotypic gender (*PG*) was equal to 0.25, to 0.24 with *ϵ*_2,*i*_ and to 0.16 with *ϵ*_3,*i*_.

## 5 Discussion

This study revealed significant phenotypic correlations between individual tree growth and reproductive traits using an original methodology combining an ecophysiological model which simulates the yearly acquisition of resource at tree level, with a hierarchical Bayesian model which specifies how the resource is allocated to growth and reproduction throughout the phenological cycle. Although similar Bayesian models have been previously developed (Buoro *et al*., 2010), this is the first time to our best knowledge that such a combined approach is used in a tree species. Considering different reproductive stage (i.e. initiation or maturation) and allowing alternatively for a driving role of NPP on the allocation scheme or not, our results shed light on the contradictory findings reported in the literature about the sign and biological meaning of the correlations between growth and reproduction (Knops *et al*., 2007).

### 5.1 Origins of the phenotypic correlations between growth and reproduction components

We detected a net increase between cone production and NPP, because trees in lower density, having more NPP tends to growth and reproduce more (Appendix S1: Figure S2). We also found a positive correlation between growth and initiation of reproductive organs: trees which invested more in reproduction had also a higher growth. This result was obtained while our model does not specify any competition between sinks in case of limiting resources. Resources appeared to be invested proportionally in bud initiation and growth with a constant fraction devoted for reproduction, which is referred as the “resource matching” hypothesis. Under this hypothesis, trees both grow and reproduce more when more resources are available (Bogdziewicz *et al*., 2020).

Then, we found a negative correlation between growth and cone survival, suggesting that these functions of growth and cone maturation compete with each other during resource allocation. Similarly, a recent study in *Fagus sylvatica* demonstrated that reproduction drives the inter-annual variability of growth and that years with high reproductive effort were also years with reduced growth (Hacket-Pain *et al*., 2018). However we cannot rule out that other processes such as pollen limitation contribute to the negative correlation observed here between growth and reproduction, as showed by Knops *et al.* (2007). Indeed, in wind pollinated species, springs with high level of precipitation usually favor growth, but can reduce the efficiency of pollen release and fertilization. In these cases, negative correlation between growth and reproduction can emerge from contrasted effects of meteorological conditions, without any direct link between growth and reproduction. But we demonstrated here that resources could also contribute to this trade-off between growth and fruit maturation and it is not only dependent to pollination.

Finally, we also found a trade-off between buds initiation and female cones survival: trees investing more resource to maturate cones initiated less reproductive buds. This result is consistent with previously published experimental data, in particular removal experiments which demonstrated a trade-off between current and future reproduction (for instance in *Pinus halepensis*, Santos-del Blanco and Climent 2014).

The three observed correlations differ according the stand densities (Fig. 5). Trees in low density tend to allocate, proportionally, more resources to growth than in high density stand, while trees in high density tend to allocate in general more resources to bud initiation and fruit maturation than in low density stand. When resources are scarce (i.e. high density stand), trees are expected to exhibit a stronger trade-off and to invest most of their resources in reproduction whereas when resources are abundant, trees are expected to invest it preferentially for growth (Lauder *et al*., 2019). Furthermore, correlations between growth and reproductive traits were only present in the high density stand, except for the last correlation between initiation and cone maturation, found in both densities. Our study also revealed that NPP is not the only driver to the different correlations. Additional drivers might be important too (such as reserves, nitrogen, or other sources of energy).

Finally, we found that *C. atlantica* trees were overall more females both at high and low competitor density, and that the phenotypic gender was relatively constant across years (expect in 2004 with a lower median value for low density plot, Fig 4).

### 5.2 Evolutionary consequences of the observed trade-offs

In the studied system, *C. atlantica* trees displayed different strategies of resource allocation: some individuals allocated resource preferentially to growth and others to cone survival; some individuals allocated resource preferentially to cone survival and other to reproductive buds initiation. These strategies were observed in an even-aged plantation where the studied individuals had the same age, experiment similar climatic conditions and also similar density of competitor within stand. As trade-offs were identified through correlations between individual random effect, they integrate the genetic and plastic inter-individual variation of traits involved in growth and reproduction. The observed trade-off cannot therefore be entirely attributed to genetic factors although we expect the plastic component of trait variation to be minimized in this plantation as compared to a natural population.

This approach could be used in natural populations, using a similar model if any proxy of spatial resource heterogeneity were available as a covariate, or a simplified model if not. Therefore, we expect that the heterogeneity of density in a natural population would also result in the coexistence of multiple allocation strategies, but along a wider range of trait values. Moreover, the heterogeneity in tree density should not only affect the amount of available resource for individuals, but also change the conditions of evolution. Finally, our study also suggests that different individual strategies are expected at the core versus the margins of natural population range (Sexton *et al*., 2009). Indeed, we showed that trees at low-density grew more than individuals at high-density, which affected the slope (although not the sign) of the correlation between growth and reproduction. But slopes of higher values where found for low density stand. Such differences in resources acquisition could also occur across time and ontogenic stages (Thomas, 2011; Barringer *et al*., 2013) and may affect the capacity of migration and adaptation (Aitken *et al*., 2008).

### 5.3 A new approach for modelling resource acquisition and allocation

The main novelty of our approach is to combine the ecophysiological model CASTANEA to simulate resource acquisition with a hierarchical Bayesian model to account explicitly for resource allocation through latent variables (e.g. *IB*_*i,t*_, *PG*_*i,t*_). CASTANEA allowed us to obtain and validate proxys of the available resource, through the carbon Net Primary Production (NPP). The model correctly reproduced the variation in the level of resource between plots at different competitor densities. Higher NPP per individual was simulated for the low density stand, consistent with reduced competition. This positive effect of reduced competition on resource acquisition at tree level was also observed with tree growth data (Guillemot *et al*., 2015) and simulated with another version of the CASTANEA model (Guillemot *et al*., 2014). Based on tree ecophysiology, climate and soil conditions, variations in the levels of resource were also simulated among years. In particular, lower resources were simulated during the year 2003 due to higher stress. Drought stress is known to directly impact ecosystem productivity (Ciais *et al*., 2005) and its duration determines the growth of Mediterranean trees (Linares *et al*., 2013; Lempereur *et al*., 2015).

A second advantage of our approach is the use of latent variables in the Bayesian model, which allowed us to explicitly model resource allocation (including the phenotypic gender) over the two-years reproductive cycle. This Bayesian model revealed the existence of trade-offs that could not observed based on raw data (Fig. 3), if we look at cone production and tree growth. Many perennial species have a similar reproductive cycle over two years, with bud initiation in the first year, followed by maturation in the second year. The hierarchical Bayesian model developed in this study could thus be applied to other species, using direct measurements of resource proxys (e.g. photosynthesis), or resources estimates obtained from an ecophysiological model (for example the CASTANEA is currently calibrated for twelve European trees species). Our model can also be used in species with a shorter reproductive cycle, where female fruit mortality would occur the same year of it initiation.

Another advantage is the use of two different resource allocation scheme in the same equation (i.e. use of *Y*) to identify correlation. It was possible to test if individual random effects were strictly based on NPP resource or other kinds of resources or processes. We found that for all iterations that model 2, with individual random effects outside the slope, was the best, meaning that allocation to growth and reproduction are the result of not only NPP. Deeper exploration is needed to investigate what are the main limiting energies responsible of these trade-offs.

### 5.4 Towards a synthesis between models of resource allocation to reproduction

With the Bayesian model, it was possible to test two potential models with a constrain on the resource allocation and no constrain and allowed us to identify inter-individual correlation. We found that trade-offs emerged from the estimation of the correlations between individual random effect without the inclusion of any constraints on resource allocation. This is not the case of other models that impose detailed on mechanisms driving resource acquisition and allocation, such as Dynamic Energy Budget model (DEB) or Resource Budget model (RBM). In DEB models, reserves used for growth cannot be used to increase reproduction, as direct competition occurs only among reserves used for building structure and not with those used for paying maintenance costs (Kooijman, 2009). Hence, the fraction of reserves used for growth and reproduction is fixed. In RBM models, the resource is allocated to reproduction only when resource level exceeds a threshold (Isagi *et al*., 1997). Indeed, RBM models consider that plants cannot have high fruit production during several years due to resource depletion (Crone and Rapp, 2014). Our results support the modeling choice used in DEB and RBM models where they include allocation to reproduction. But modeling explicitly the variation of resource partitioning between growth and reproduction across the phenological cycle could improve these models.

We also found here that accounting for the reproductive phenology, which determines the timing of resource allocation, can improve the estimation of trade-off between growth and reproductive sinks, and can highlight the idiosyncratic correlation patterns observed in trees so far. Indeed, in the study system, the sign of the correlation between growth and reproduction differs depending of the reproductive stage (initiation or maturation). But with shorter reproductive cycle, we could expect different results for the relation between growth and fruit initiation, because plants may invest resources for initiation and maturation during the same year.

## 6 Conclusion

Combining a simulation, process-based model with a Bayesian statistical estimation model can be a fruitful approach to investigate the potential trade-off between reproduction and growth. Process-based models integrate species ecology through their ecophysiological characteristics, and allow to explicitly model the effects of environmental stresses on reproduction, and in particular on reproductive phenology. However, the processes involved in tree reproduction are still not well understood and most models assume that a constant fraction of resource is allocated to reproduction (Vacchiano *et al*., 2018). Further improvements of the resource allocation component in these models are needed to understand how tree growth and reproduction jointly respond to climate change.

## 7 Acknowledgements

We thank F. Jean, W. Brunetto, O. Gilg, D. Vauthier and F. Rei for data collection. This work is part of the PhD thesis of the first author, funded by the Provence-Alpes-Cote-d’Azur region and the INRA institute. We acknowledge E. Klein, C. Perret and T. Caignard for useful comments, F. Jean and A. Chalon for providing photos for the Figure 2. We also thank reviewers for their useful comments on previous versions. This work was supported by the metaprogramme Adaptation of Agriculture and Forests to Climate Change (AAFCC) of the French National Institute for Agricultural Research (INRA) and by the ANR-13-ADAP-0006 project MeCC.

## 8 Author Contribution

V.J., F.L. & H.D. initiated the idea. F.L & F.C. provided experimental data. H.D. and V.J. developed the ecophysiological model for *C. atlantica*. J.P. & E.W. designed the Bayesian model with advice from all authors. V.J., E.W. and J.P. ran simulation. V.J. started the redaction in consultation with F.L., H.D. and S.O-M. All authors contributed critically to the drafts and gave final approval for publication.

## 9 Data accessibility statement

Data used, CASTANEA model input simulation and hierarchical Bayesian model code are available from the Zenodo repository <https://doi.org/10.5281/zenodo.4433892>. The CASTANEA model is an open-source software available on the CAPSIS plateform: http://capsis.cirad.fr/capsis/models

