## supporting information for "A hierarchical Bayesian model to investigate trade-offs between growth and reproduction in a long-lived plant"

**Appendix S1**
**Supplementary sections**
**Section S1**: Description of the CASTANEA model, calibration and validation for *Cedrus Atlantica*
**Section S2**: Bayesian Model implementation and analysis

**Supplementary tables and figure**
**Table S1**: Prior distribution definition
**Table S2**: Posterior predictive check of the hierarchical Bayesian model
**Table S3**: Summary of posterior distribution parameters
**Figure S1**: relation between traits (number of cones or growth) and NPP.

### 1 Supplementary Sections

#### 1.1 Overview of the CASTANEA model

CASTANEA is an ecophysiologically-based model simulating carbon and water fluxes in forest ecosystems with no spatial-explicit representation of trees (Dufrêne et al., 2005). The original CASTANEA model considers an average tree, represented by six functional compartments: leaves, branches, stem, carbon reserves (i.e Non Structural Carbon Content), fine roots and coarse roots. Tree canopy is divided into five layers of leaves.

The physiological processes included in the model are radiation transfer, photosynthesis, maintenance and growth respiration, carbon allocation, water interception, transpiration and soil evaporation. Three different radiative balances are half-hourly simulated, in the Photosynthetically Active Radiation (PAR, 400–700 nm), in the Near-Infrared (NIR, 700–2500 nm), and in the thermal infrared for each canopy layer using an adaptation of SAIL model (Verhoef, 1984, 1985). Photosynthesis is half-hourly simulated for each canopy layer using the model of Farquhar et al. (1980) and analytically coupled to the stomatal conductance model proposed by Ball et al. (1987). Maintenance respiration is simulated as proportional to the nitrogen content for each considered organs (Ryan, 1991). Growth respiration is simulated from growth increment combined with a construction cost specific to organ's types De Vries et al. (1974). Transpiration is hourly simulated based on Monteith (1965) equations. Soil evaporation is simulated in the same way as canopy transpiration using Penman–Monteith equation. The dynamics of soil water content is then simulated each days. Soil drought drives stomata closure via a linear decrease in the slope of the stomatal conductance model (Ball et al., 1987), when relative soil water content is under 40% of field capacity (Sala and Tenhunen, 1996; Granier et al., 2000). In the carbon allocation sub-model, allocation coefficients between compartments (i.e. fine roots, coarse roots, stem, branches, leaves and reserves) are simulated each days depending on the sink and the phenological constraints (Davi et al., 2009; Davi and Cailleret, 2017). Budburst date was simulated with a one-phase model, which describes the cumulative effect of forcing temperatures on bud development during the ecodormancy phase (Dufrêne et al., 2005).

#### 1.2 CASTANEA parameterization

The CASTANEA model was originally developed and validated from organ to stand-scale for *Fagus sylvatica* in Davi et al. (2005), *Pinus sylvestris*, *Pinus pinaster* in Davi et al. (2006), *Picea abies* in Delpierre et al. (2012) and *Abies alba* in Davi and Cailleret (2017) with forest carbon fluxes (e.g. Gross Primary Productivity and Ecosystem Respiration) or with tree growth increment (ring width). In this study, we used data from literature and *in situ* to parameterize CASTANEA for *Cedrus atlantica* (Table S1- 1), and validated it with ring width data.

For each density plot, we accounted for the soil and stand characteristics. For the soil we included parameters describing soil height, percentage of stones, clay/sand percentage in top and deep soil, humidity at wiling point and at field capacity. We simulated a total of 71 individuals, accounting for their DBH, tree volume and leaf area index differences.

The climate variables used as model input are daily precipitations (in $\mathrm{mm}$), minimal, maximal and average temperature (in $\mathrm{T^{\circ}C}$), global radiation (in $\mathrm{mJ/d}$), relative humidity (in$\%$) and wind speed (in $\mathrm{m/s}$). We used here local climatic variables measurements from 1999 to 2005, from a climate station located at 1.96 km from the study site. Local daily climate were estimated from 1989 to 2015 with a long-term national meteorological data and down-scaled with statistical regressions (Quintana-Seguí et al., 2008).

In the model we also accounted for thinning intensity, where a percentage of trees were cut at different years (clearcut during the year 1991 and 2002 for the density plot 250, and in 1991, 1998 and 2014 for the density plot 1200). Thinning reduces the stand leaf area index and the stand density, thus increasing the resources available for the remaining trees.

All files used for model simulation are presented at <`https://doi.org/10.5281/zenodo.4433892`>. We reported the inventory file which contains all site and individual characteristics with different model options (allocation rules, phenology process), the climate data file used as climate forcing variable and the forestry files where it is indicated year and percentage of clearcut. We used here the CASTANEA model available from the community modelling platform CAPSIS (version 15325, Dufour-Kowalski et al. 2012).

#### 1.3 CASTANEA model validation and simulation design

Measurements and simulations were done for each individuals from the two density plot presented in the main text, and from 1989 to 2015. We used diameter increment measurement for 40 individual trees in the high-density plot and 31 individual trees in the low-density plot. We measured the diameter at 1.3 meters ($DBH_{i,t}$) for each individual tree $i$ and each year $t$ and converted it into ring width increment ($rw_{i,t}$, in mm). $rw_{i,t}$ was computed as: $rw_{i,t} = (DBH_{i,t}/\pi/2) - (DBH_{i,t-1}/\pi/2)$. We then compared simulated ring width from the CASTANEA model with ring width increment observation. We assessed the ability to predict ring width increment with three metrics; the *Root Mean Squared Error* (RMSE), the *Coefficient of determination* ($R^2$) and the *percent bias* (PB). The RMSE gives the standard deviation of model prediction error. The $R^2$ give the proportion of variance of one variable that is predictable from the other variable, ranging between 0 to 1 if simulation is equal to observations. The PB measures the average trend of simulation to be higher or smaller than observations, with an optimal

| Abbreviation | Variable (units) | value |
|---|---|---|
| $CR_{leaves}$ | Leaf construction cost ($gC.gC^{-1}$) | 1.32 |
| $CR_{coarseroots}$ | Coarse roots construction cost ($gC.gC^{-1}$) | 1.364 |
| $CR_{fineroots}$ | Fine roots construction cost ($gC.gC^{-1}$) | 1.28 |
| $CR_{wood}$ | Wood construction cost ($gC.gC^{-1}$) | 1.364 |
| $TGSS$ | Initial Non-Structural Carbohydrates concentration ([NSC]) ($gC.gC^{-1}$) | 0.2 |
| $N_{leaves}$ | Nitrogen in leaves (%) | 1.52 |
| $\Psi_{wood}$ | Predawn potential for wood growth cessation (MPa) | -1 |
| $F_{rootstoleaves}$ | Ratio between fines roots and leaves biomass | 0.3 |
| $LA$ | Leaf area | 0.0000104 |
| $LMA_{sunmax}$ | Leaf Mass per Area of sun leaves | 245 |
| $\alpha_L$ | Leaf angle inside the canopy (radians) | 40 |
| $\alpha_b$ | Branches angle inside the canopy (radians) | 8.7 |
| $CR_1$ | Slope of the crown area to dbh-relation | 0.08035 |
| $CR_2$ | Intercept of the crown area to dbh-relation | 0.6281 |
| $aGF$ | Slope of the height-dbh relationship | 4.356199 |
| $bGF$ | Power coefficient of the height-dbh relationship | 0.3646 |
| $\phi$ | Form coefficient of stem | 0.41‡ |
| $\rho_{wood}$ | Wood density ($kg.m^{-3}$) | 545.2§ |
| $CF$ | canopy clumping coefficient | 0.46 |
| $\rho_{wPIR}$ | wood reflectance for PIR radiation | 0.49 |
| $\rho_{wPAR}$ | wood reflectance for PAR radiation | 0.15 |
| $\rho_{lPIR}$ | leaf reflectance for PIR radiation | 0.29 |
| $\tau_{lPIR}$ | leaf transmittance for PIR radiation | 0.228 |
| $\rho_{lPAR}$ | leaf reflectance for PAR radiation | 0.096 |
| $\tau_{lPAR}$ | leaf transmittance for PAR radiation | 0.047 |
| $g_0$ | intercept of ball and berry relation | 0.000492 |
| $g_1$ | slope of ball and berry relation | 10.5† |
| $P_{50}$ | Water potential inducing 50% loss of conductivity (MPa) | -5.482 |
| $\alpha_{Na}$ | Dependency between $V_{Cmax}$ and leaf nitrogen ($\mu molCO^2gN^{-1}s^{-1}$) | 10.47† |
| TSUMBB | Critical value of state of forcing (from dormancy to active period) | 365 |
| $woodStop$ | Critical value of state of forcing from date of onset to end of wood growth | 600 |
| $T_{min_{EB}}$ | minimal temperature below which frost has an effect on young buds (°C) | -4.1 |
| $Cohorte_{leaves}$ | maximum needle or leaves lifespan | 3 |
| $pleafMin$ | minimal leaf potential | -2.9 |
| aRDI | intercept parameter used for relative density index (RDI) | 12.5 |
| bRDI | slope parameter used for relative density index (RDI) | -1.651 |

Table S1- 1: List of species-specific parameters of CASTANEA for *Cedrus atlantica* from *in situ* measurements and literature. † parameter adjusted from Ladjal et al. (2007), ‡ from Courbet (1991), § from Hapla et al. (2000).

value at 0, overestimation bias with positive values and underestimation with negative values. All these metrics were computed from the average values of observations or simulations over all years and for both densities.

Once the model validated, the aim of the simulation was to simulate individual tree productivity, accounting for their individual characteristics in terms of DBH and volume, and thinning intensity between the two plots. To do so, we converted the Net Primary Productivity (NPP, originally in gC.m$^{-2}$.year$^{-1}$) into individual and annual NPP based on tree crown projection with the following equation:

$$NPP_{i,t} = NPP_t \times \frac{\pi \times (CR_1 \times DBH + CR_2)^2}{RDI} \tag{1}$$

With $DBH$ for tree diameter, $CR_1$ for the slope of the crown area to dbh-relation, $CR_2$ the intercept of the crown area to $DBH$ relationship, $RDI$ is the relative density index and $NPP_t$ is the NPP per m$^{-2}$ of soil. Variation of NPP is presented in the main text for both density plots (see main text, Fig. 4).

The RDI is obtained with the equation based on the study of Charru et al. (2012):

$$RDI = \frac{N}{\exp aRDI + bRDI * \log_{Dg}} \tag{2}$$

With $aRDI$ and $bRDI$ are respectively intercept and slope parameter, $N$ is the number of trees per ha, and $Dg$ is the log quadratic mean diameter (m).

#### 1.4 Climate data

Local climatic variables were measured from 1999 to 2005 using a meteorological station (Hobos ProV2 micro-loggers) located at 1.96 km from the study site. However, we needed a longer period to calibrate and validate the model CASTANEA. Local daily climate was obtained from long-term national meteorological data from 1989 to 2015 and down-scaled using statistical regression on local climatic measurements (Quintana-Seguí et al., 2008). We used precipitations (in $\mathrm{mm}$), minimal/maximal/average temperature (in T°C), global radiation (in $\mathrm{mJ\,d^{-1}}$), relative humidity (in $\%$) and wind speed (in $\mathrm{m\,s^{-1}}$) as forcing variables for CASTANEA.

#### 1.5 Validation results

Ring widths simulated by CASTANEA over the period from 1989 to 2015 were well correlated with observed ring widths with a $R^2$ of 0.44 and 0.20 for high and low density plot, respectively (Fig.S1- 1). A overestimation was observed for the high density plot and underestimation for the low density plot. Two peaks of ring width were observed for 2011 and 2014 possibly due to

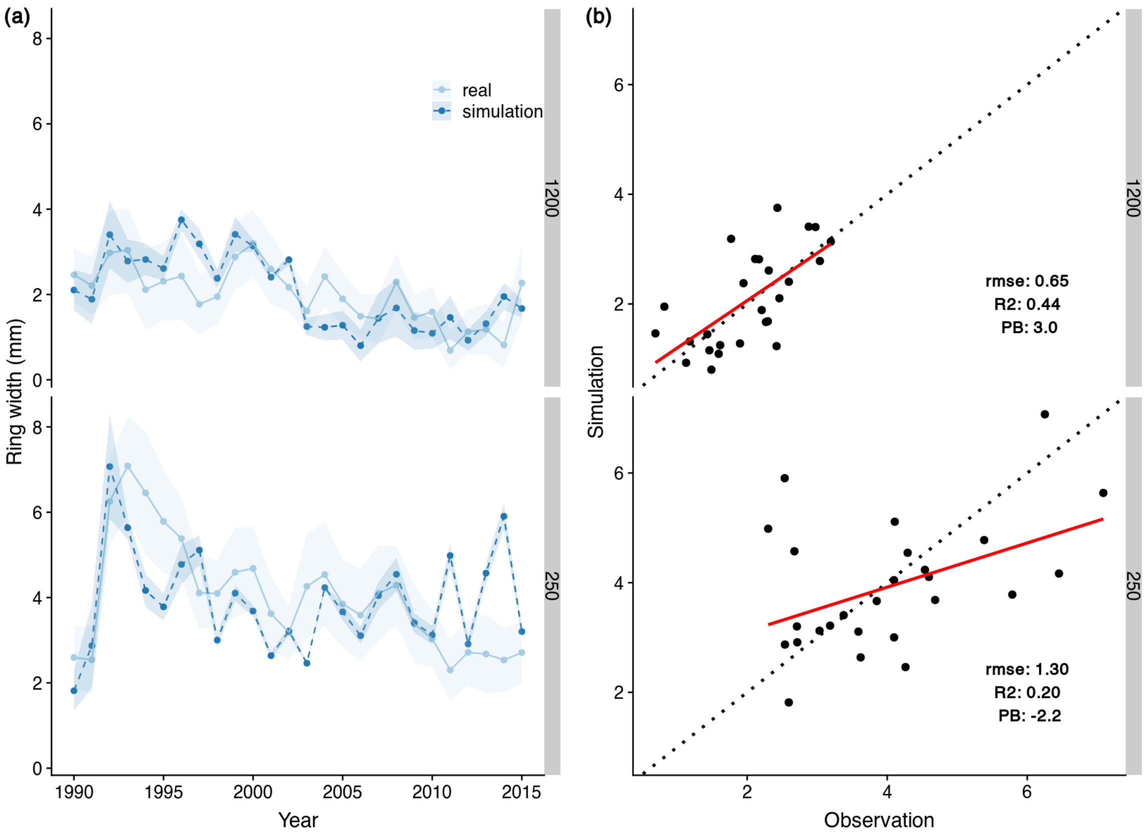


Figure S1- 1: Comparison between ring width observation and simulation from 1989 to 2015 and for both density plots. (a) Ring width observation is presented in pale blue and ring width simulations in darker blue. Blue pale line correspond to mean observations, blue dashed line for mean simulation, with their standard deviation in both envelope. (b) Black dotted line is the 1:1 line with red line is the linear regression between simulated values of ring width and observations. Goodness-of-fit of the model is present with the $RMSE$, $R^2$ and $PB$.

an advance of budburst and a low hydraulic stress which favor tree growth (not presented here). We presented NPP used for the hierachical Bayesian model in the main text.

#### 2 Bayesian Model implementation and analysis

The prior distributions of the model parameters are reported in Appendix S1: Table S1. We performed the estimation of parameters with the `JAGS` software (v4.3.0, Plummer 2003) in the `R` environment (v3.6.3, R Core Team 2018). We ran three parallel chains with 100000 iterations, a burn-in of 100000 and a thinning rate at 100. In the result section, we presented posterior median along with the 95% credible interval of parameters. We conducted posterior predictive checks to evaluate the accuracy of our model in fitting the data. To do so, we simulated replicated data under the fitted model and compared them with the observed data. We then calculated a Bayesian p-value for quantitative variables, $BAI_{i,t}^{\text{obs}}$ and $FC_{i,t}^{\text{obs}}$. A value near 0.5 indicates that the model correctly fits the data (Gelman et al., 1996). For male cone notation ($IMC_{i,t}^{\text{obs}}$), which only takes 0 and 1 values, we used the Brier score to determine its accuracy. This score varies between 0 and 1 with a score below 0.25 indicating that the model predicts better than expected by chance.

We also checked how $\epsilon$ terms are influenced by the phenotypic gender ($PG_{i,t}$) with linear regression in each iteration and reported probabilities to obtain $P < 0.05$. We performed linear regressions with the `stats` package (R Core Team, 2018). We reported posterior predictive checks, parameters values with median posterior and credible interval in the main text and in Appendix S1: Table S2 and Table S3.

### 3 Supplementary Tables and Figure

| **Parameters** | **Prior distribution** |
|---|---|
| $\gamma_d$ | $\gamma_d \sim \mathcal{N}\,(0, 0.001)$ |
| $\beta_{1,d}$ | $\beta_{1,d} \sim \mathcal{N}\,(0, 0.001)$ |
| $\beta_{2,d}$ | $\beta_{2,d} \sim \mathcal{N}\,(0, 0.001)$ |
| $\beta_0$ | -10 |
| $p_X$ | $p_X \sim \mathcal{U}\,(0, 1)$ |
| $p_Y$ | $p_Y \sim \mathcal{U}\,(0, 1)$ |
| $\sigma_{BAI}$ | $\sigma_{BAI} = 1/\sigma_d^2$ with $\sigma_d \sim \mathcal{U}\,(0, 10)$ |
| $\sigma_{IMC}$ | $\sigma_{IMC} = 1/\sigma_r^2$ with $\sigma_r \sim \mathcal{U}\,(0, 50)$ |
| $\bar{PG}$ | $\bar{PG} \sim \mathcal{N}\,(0, 0.001)$ |
| $\sigma_{PG}$ | $\sigma_{PG} = 1/\sigma^2$ with $\sigma \sim \mathcal{U}\,(0, 10)$ |
| $\Sigma$ | $\Sigma \sim \mathcal{W}\,(Id3, 4)$ |

Table S1: Description of prior distribution. $\mathcal{N}$ for $Normal$ distribution, $\mathcal{U}$ for $Uniform$ distribution and $\mathcal{W}$ for the inverse $Whishart$ distribution

| **Observed variable** | **Bayesian posterior p-value** |
|---|---|
| Growth ($BAI_{i,t}^{\text{obs}}$) | 0.18 |
| Female cone number ($FC_{i,t}^{\text{obs}}$) | 0.84 |

| **Observed variable** | **Brier score** |
|---|---|
| Male notation ($IMC_{i,t}^{\text{obs}}$) | |
| $note$ 0 | 0.273 [0.265; 0.282] |
| $note$ 1 | 0.080 [0.080; 0.081] |
| $note$ 2 | 0.129 [0.128; 0.129] |
| $note$ 3 | 0.090 [0.089; 0.090] |
| $note$ 4 | 0.211 [0.202; 0.220] |

Table S2: Summary posterior predictive check. For growth and female cone number we reported Bayesian $P$-values with a value at 0.5 for a perfect fit. For male notation, we reported the Brier score with median and 95% CI, with value closed to 0 for correct fit.

| Parameters | Median and CI values |
|---|---|
| $PG$ | -1.42 [-2.31; -0.75] |
| $\sigma$ | 2.13 [1.63; 2.90] |
| $\sigma_r$ | 47.79 [42.68; 49.89] |
| $\sigma_d$ | 7.39 [6.89; 7.94] |
| $p_Y$ | 0.29 [0.01; 0.85] |
| $\gamma_{1200}$ | 43.6 [39.3; 48.1] |
| $\gamma_{250}$ | 46.2 [42.0; 50.0] |
| $\beta_{1,1200}$ | 1.98 [0.41; 3.39] |
| $\beta_{1,250}$ | 1.03 [0.67; 1.42] |
| $\beta_{2,1200}$ | 30.1 [26.7; 35.3] |
| $\beta_{2,250}$ | 26.6 [23.0; 31.2] |

Table S3: Summary of posterior distribution (medians and credible intervals) for parameters used in our model.

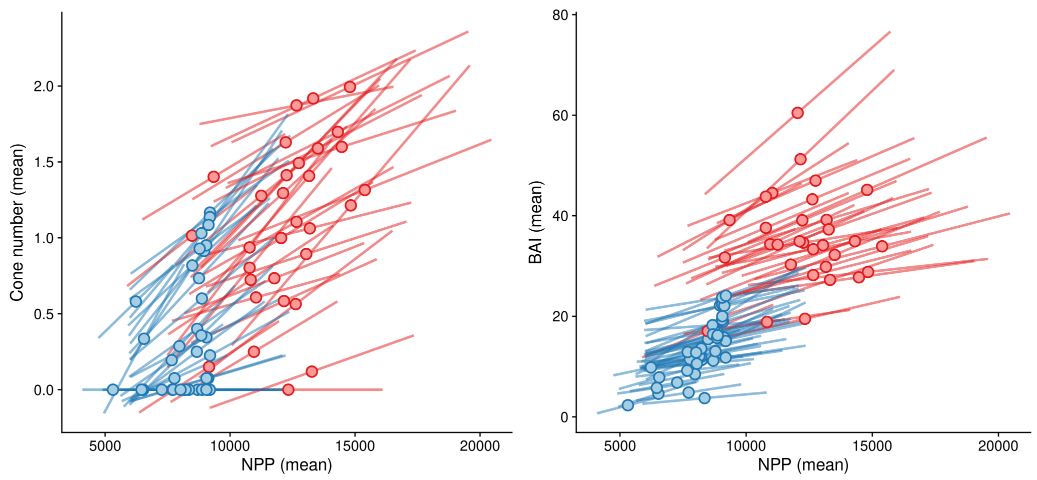


Figure S1: Relation between cone production and simulated NPP from CASTANEA (left). Relation between tree growth (BAI) and NPP simulated (right). Each dots is mean value with it standard deviation, with blue for individuals in high density (1200) and red for low density stand (250). For cone production, we computed log10(1+cone number) due to null observations.
